# A joint reaction coordinate for computing the free energy landscape of pore nucleation and pore expansion in lipid membranes

**DOI:** 10.1101/2020.09.23.309898

**Authors:** Jochen S. Hub

## Abstract

Topological transitions of membranes, such as pore formation or membrane fusion, play key roles in biology, biotechnology, and in medical applications. Calculating the related free energy landscapes has been complicated by the fact that such processes involve a sequence of transitions along highly distinct directions in conformational space, making it difficult to define good reaction coordinates (RCs) for the overall process. In this study, we present a new RC capable of driving both pore nucleation and pore expansion in lipid membranes. The potential of mean force (PMF) along the RC computed with molecular dynamics (MD) simulations provides a comprehensive view on the free-energy landscape of pore formation, including a barrier for pore nucleation, the size, free energy, and metastability of the open pore, and the energetic cost for further pore expansion against the line tension of the pore rim. We illustrate the RC by quantifying the effects (i) of simulation system size and (ii) of the addition of dimethyl sulfoxide (DMSO) on the free energy landscape of pore formation. PMF calculations along the RC provide mechanistic and energetic understanding of pore formation, hence they will be useful to rationalize the effects of membrane-active peptides, electric fields, and membrane composition on transmembrane pores.

## Introduction

The opening and closing of pores over lipid membranes play critical roles in various processes of biophysical, biotechnological, or pharmaceutical relevance. Many antimicrobial peptides function via the formation of transmembrane pores, thereby causing the loss of ionic concentration gradients over bacterial membranes.^1^ Pore-forming toxins are produced as virulence factors by many pathogenic bacteria. ^2^ Pore formation is a critical step during membrane fusion, as required for the infection of cells by enveloped viruses, for exocytosis, and for other forms of active transport.^3,4^ In turn, pores are required to close during endocytosis, that is, by transport of cargo into the cell via the formation of an intracellular vesicle. Pores are also formed on purpose with applications in medicine, for instance with the aim of drug delivery, gene therapy, or killing of malignant cells.^5^ For biotechnological applications, the opening of membranes pores is often induced by applying electric fields, a process termed electroporation.^6,7^ In biophysical experiments, pores were also induced by the application of osmotic stress or membrane tension.^8–10^ A quantitative understanding of such processes requires understanding of the free-energy landscape of pore formation.

Computing the free energy landscape of pore formation, for instance from molecular dynamics (MD) simulations with a biasing method such as umbrella sampling or constrained simulations, is complicated by the fact that pore formation involves two highly distinct conformational transition, here referred to as (a) pore nucleation and (b) pore expansion. ^11^ Because pore nucleation and pore expansion occur along different directions in conformational space, it has been difficult to define a unified reaction coordinate (RC) for the overall process. Namely, during nucleation, the membrane is locally thinning, followed by the penetration of water and head groups into the hydrophobic core of the membrane.^12–14^ Hence, nucleation is primarily characterized by a transition of polar atoms *perpendicular* to the membrane plane. In contrast, expansion of an established pore involves the increase of the pore radius, that is a transition of the (already established) pore rim *parallel* to the membrane plane. Notably, the expansion of large pores was successfully modeled by classical nucleation theory (CNT).^15,16^ CNT describes the pore free energy as function of the pore radius *R* as

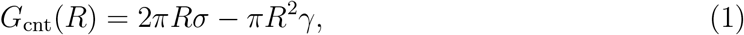

where *σ* denotes the line tension of the pore rim, and *γ* is the surface tension between membrane and water. However, because CNT breaks down at small pore radii, it is not suitable for describing the energetics during pore nucleation.^11^

Early simulation studies of pore formation were carried out under non-equilibrium conditions, where the pore was induced by applying an external perturbation, such as an electric field or mechanical stress.^17–21^ From such non-equilibrium simulations, it is difficult to obtain the free energy cost of pore formation. To obtain the potential of mean force (PMF) of pore formation, some studies used the pore radius *R* as RC.^22,23^ Because *R* is not suitable for following the early steps of pore formation during the nucleation process, several alternative RCs have been proposed, including a RC that steers the lipids collectively in lateral direction away from the pore center,^24^ a lipid flip–flop coordinate, steering a single lipid head group towards the membrane core,^13,25–27^ or the mean solvent density within a membrane-spanning cylinder.^28^ Complementary, a theory for pore formation based on continuum elasticity was recently proposed, which quantified the pore free energy in terms of the pore radius together with the thickness of a hydrophobic pore rim.^29,30^

In a systematic comparison of RCs of pore formation, we found that pulling along established RCs for pore nucleation efficiently triggered pore formation. However, if the open pore is metastable (long-living), pulling along these RCs in reverse direction did not reliably close an established pore, primarily because the continuous hydrogen bond network along the transmembrane pore was not reliably disrupted by pulling along these RCs. ^31^ Such different pathways during forward and reverse pulling may lead to severe hysteresis problems during PMF calculations. The problems prompted us to develop a RC that probes the degree of connectivity of a polar transmembrane defect, here referred to as “chain coordinate” *ξ*_ch_, which is briefly described in the Methods. We found that pulling along *ξ*_ch_ allows for reliable pore opening *and* closing, thereby avoiding any hysteresis problems during PMF calculations.^11,32^ Using the chain coordinate, we resolved for the first time a nucleation barrier and, thereby, the metastability of the open pore in certain membranes, which had been proposed forty years ago to rationalize memory effects in electrophysiology experiments.^6^ An alternative biasing potential for efficient pore opening and closing, coined “gizmo”, was recently proposed based on a set of repellant beads.^14^ Here, a linear, membrane-spanning arrangement of beads was used to drive pore opening, and a ring-shaped arrangement of beads was used to squeeze a transmembrane polar defect to drive pore closing. In the same study, a careful committor analysis showed that the penetration of head groups into the membrane core, and not merely of water, is critical for crossing the transition state of pore nucleation.

While the chain coordinate *ξ*_ch_ has been designed to follow pore nucleation and to resolve a nucleation barrier (if present), it is not suitable to follow pore expansion because all pores with larger radii are projected onto *ξ*_ch_ = 1, characterizing a fully formed transmembrane defect. Consequently, PMFs along *ξ*_ch_ do not clearly resolve the free energy minimum and the size of an open, metastable pore, and they do not provide the free energy cost for further pore expansion against the line tension of the pore rim. In this study, a new RC is introduced that combines nucleation along *ξ*_ch_ with expansion along the pore radius. We show that a single PMF along the new RC reveals the complete free energy landscape reflecting the unperturbed membrane, pore nucleation, a metastable pore, and further pore expansion. We illustrate the RC by analyzing the effects of membrane size and of dimethyl sulfoxide (DMSO) on the free energy landscape of pores.

## Methods

### Definition of the reaction coordinate (RC)

The new RC is defined as a combination of two coordinates. First, we quantify the state of pore nucleation with the chain coordinate *ξ*_ch_(**r**) defined in Ref. 32, in which it was simply denoted by “*ξ*”. The symbol **r** denotes the coordinates of all atoms of the system. *ξ*_ch_(**r**) was designed to quantify the connectivity of a polar transmembrane defect. In brief, *ξ*_ch_(**r**) is defined via a transmembrane cylinder that is decomposed into *N_s_* slices with a thickness of typically 1Å, as illustrated in Fig. 1A. *ξ*_ch_(**r**) is approximately given by the *fraction* of slices that are filled by polar heavy atoms (Fig. 1A, blue shaded slices). As polar atoms contributing to *ξ*_ch_, we typically used oxygen atoms of water and lipid phosphate groups. Hence, both water and head groups contribute to the degree of connectivity that is quantified by *ξ*_ch_. Upon pulling the system along *ξ*_ch_, the slices are filled one-by-one, thereby enforcing the formation of a chain of hydrogen-bonded polar atoms – hence the notion “chain coordinate”. The length of the cylinder is chosen such that *ξ*_ch_ takes a value of ~0.2 for the flat, unperturbed membrane, indicating that 20% of the cylinder slices located at the two head group regions are filled by polar atoms. A value *ξ*_ch_ ≈ 1 indicates a fully formed pore. A thin continuous water needle, flanked by indented head groups, is typically given by *ξ*_ch_(**r**) ≈ 0.85, corresponding to the transition state of pore formation in membranes that exhibit a pore nucleation barrier. Because all larger pores with increased radii are projected onto *ξ*_ch_ = 1, the chain coordinate is inappropriate for studying pore expansion. Instead, pulling along *ξ*_ch_ may efficiently open or close transmembrane defects.

**Figure 1:**
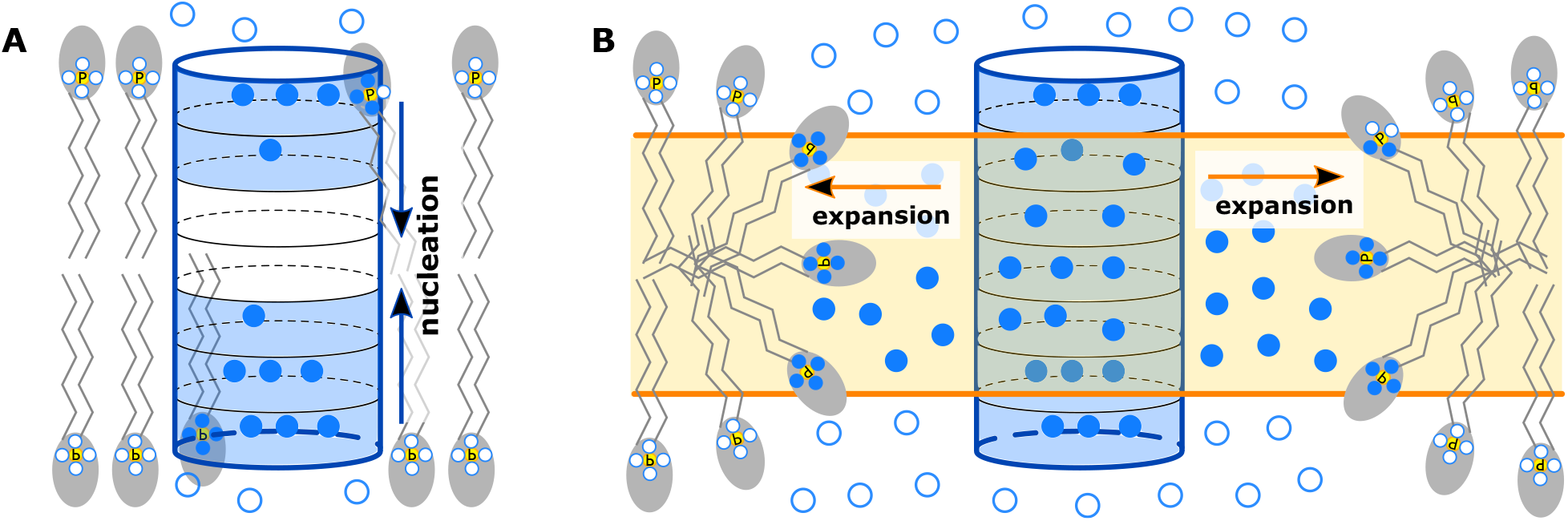
Illustration of the reaction coordinate. (A) The chain coordinate *ξ*_ch_ quantifies the connectivity of a polar defect as the fraction of polar-occupied slices of a transmembrane cylinder. Polar-occupied slices are highlighted by blue shading. Only polar heavy atoms within the cylinder contribute to *ξ*_ch_ (solid blue circles), but not polar atoms outside the cylinder (empty blue circles). We considered oxygen atoms of water and phosphate groups as polar atoms contributing to the reaction coordinate. (B) After a continuous polar defect has formed (*ξ*_ch_ ≈ 1), the pore expands along the radius of the pore, quantified via the number of polar heavy atoms within a horizontal slab at the membrane center (blue solid circles inside the orange shading).

Second, we quantify the state of pore expansion via the approximate radius of the pore *R*(**r**), which we define via the number of polar heavy atoms *n_P_* (**r**) within a horizontal layer of thickness *D* at the center of the membrane (parallel to the membrane plane, see Fig. 1B). Because the defect takes approximately the shape of a cylinder, we define *R*(**r**) as

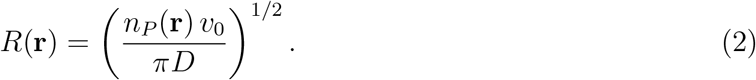

We used *D* = 1 nm in this work. The symbol *v*_0_ denotes the volume per polar atom, here taken from the molecular volume of water, *v*_0_ = 29.96 Å^3^. Hence, pore expansion is quantified via the number of polar atom *n_P_* inside the hydrophobic membrane core; we merely translate the latter quantity into an approximate pore radius to obtain a more intuitive interpretation of the RC. Similar to previous studies,^28,32^ *n_P_* was computed using a differentiable indicator function that takes unity inside the horizontal layer, and zero outside, as follows:

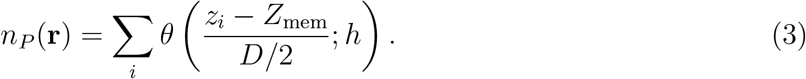

Here, *θ*(*x*; *h*) is a differentiable step function that is zero for |*x*| > 1 + *h*, and unity for |*x*| < 1 − *h*, where *h* = 0.1. For the definition of *θ*(*x*; *h*) and its derivative we refer to previous work.^32^ *Z*_mem_ denotes the center of mass of a reference group, here defined by all heavy atoms of the lipid tails.

We combine these two coordinates for pore nucleation (*ξ*_ch_(**r**)) and pore expansion (*R*(**r**)) into a single RC as follows:

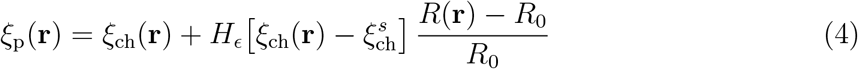

Here, *H_ϵ_*(*x*) denotes a smoothed, differentiable Heaviside step function switching between 0 and 1 in the interval [−*ϵ, ϵ*], implemented using 3^rd^-order polynomials as follows:

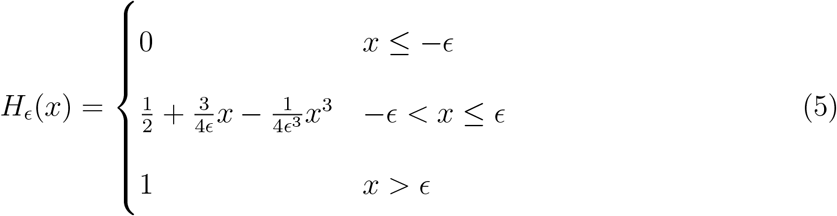

Hence, 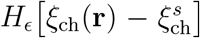 switches gradually between zero and one as the chain coordinate *ξ*_ch_(**r**) passes a switching value 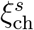. We used 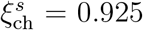, characterizing a continuous polar defect, and we used *ϵ* = 0.05. The parameter *R*_0_ serves as a normalization constant with two purposes: first, it renders the second expression on the right hand side of Eq. 4 unitless; second, by setting *R*_0_ to the value that *R*(**r**) takes where 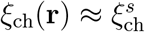, the term [*R*(**r**)−*R*_0_]*/R*_0_ takes zero in the switching region between pore nucleation and pore expansion, and then grows to finite values as the pore expands to larger radii.

Taken together, the proposed RC *ξ*_p_(**r**) quantifies the radius of the pore in units *R*_0_. The range 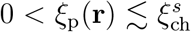 follows the *nucleation* of the pore, up to the formation of a thin pore with a radius of approximately *R*_0_. Larger values of *ξ*_p_(**r**) follow the *expansion* a fully formed defect with a radius of approximately *ξ*_p_(**r**)*R*_0_.

### Choosing the parameter *R_0_*

The parameter *R*_0_ does not change the overall shape of the free energy landscape, as different values of *R*_0_ merely shift and stretch the RC. However, to make sure that *ξ*_p_(**r**) quantifies the pore radius in units of *R*_0_, we choose *R*_0_ as the average value of *R*(**r**) in the switch region, i.e.,

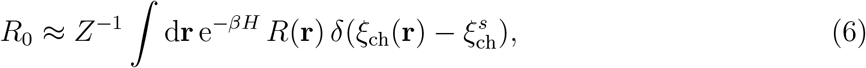

where *Z* is the partition function, *H* the Hamiltonian, *β* the inverse temperature, and *δ* the Dirac delta function. Specifying *R*_0_ via Eq. 6 would require a simulation of pore formation prior to the production simulations.

However, we found that a reasonable estimate for *R*_0_ can be obtained as follows, thereby avoiding the need of such a prior simulation. Let the chain coordinate *ξ*_ch_ be defined by a cylinder with height *N_s_d_s_*, composed of *N_s_* slices with thickness *d_s_*. The thickness *D* of the central layer is smaller than the height of the cylinder, *D < N_s_d_s_*; hence *N_s,i_* = *D/d_s_* and *N_s,o_* = *N_s_* − *D/d_s_* cylinder slices are located inside and outside of the central layer, respectively. Then, we estimated the number of polar atoms in the central layer (and thereby the value of *R*(**r**)) that leads to 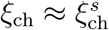. To this end, we assume that the *N_s,o_* slices outside of the central layer are fully occupied by polar atoms before a continuous polar column is formed inside the layer, yielding *ξ*_ch_ = *N_s,o_/N_s_* (the fraction of fully occupied slices). Next, we randomly add polar atoms to the *N_s,i_* slices inside the central layer, until 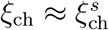. This procedure is repeated 10000 times the beginning of the simulations. Finally, the average number of polar atoms 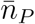 added to central layer was used to estimate *R*_0_ via Eq. 2.

We found that this estimate is in good agreement with the average *R*(**r**) in simulations with 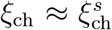. For instance, during umbrella sampling simulations with 128 DMPC lipids (*N_s_* = 26, *D* = 1 nm, *d_s_* = 0.1 nm, see below), in which the average value of the switch function was 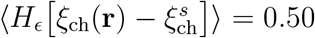 (indicating 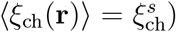, we found 〈*R*(**r**)〉 = 0.448, close to the value 0.443 for *R*_0_ obtained with the simple procedure presented above. As such, *R*_0_ is not a free parameter but is specified at the beginning of each simulation based on the parameters *N_s_*, *D*, *d_s_*, and 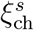.

### Implementation

The implementation of the RC *ξ*_ch_ was taken from our previous study,^32^ but merged with Gromacs 2018.8.^33^ The new RC *ξ*_p_ was implemented directly into the pull code of Gromacs 2018.8. Because the calculations of the RC and the pull forces use OpenMP parallelization, the code ran only approx. 7% slower as compared to a free simulation on a server equipped with a 6-core Intel Xeon E-2136 and an Nvidia GeForce GTX 1070Ti graphics card.

### Simulation setup and parameters

The membrane systems were set up with the MemGen webserver (http://memgen.uni-goettingen.de).^34^ Equilibrium simulations were carried out with Gromacs 2019.6, and simulations using the new RC with an in-house modification of Gromacs 2018.8. Lipids were modelled with the force field by Berger *et al.*,^35^ and the SPC water model was used.^36^ Parameters for DMSO were taken from Ref. 37. Bonds and angles of water were constrained with the SETTLE algorithm.^38^ All other bonds were constrained with LINCS.^39^ The temperature of the simulations was controlled at 300 K using velocity rescaling and by coupling water and lipid to separate heat baths.^40^ During equilibration, the pressure was controlled at 1 bar using a semi-isotropic weak coupling scheme (*τ* = 1 ps).^41^ Although weak coupling does not yield a well-defined ensemble, we used it here owing to its numerical stability. Electrostatic interactions were calculated using the particle-mesh Ewald method. ^42,43^ Dispersion interactions and short-range repulsion were described by a Lennard-Jones (LJ) potential, which was cut off at 1 nm. An integration time step of 5 fs was applied. All systems were equilibrated until the box dimensions and potential energies were fully converged.

### Simulations of pore opening and umbrella sampling simulations

PMFs were computed using umbrella sampling simulations.^44^ Starting frames for umbrella sampling simulations were taken from constant-velocity pulling simulations either along *ξ*_ch_ or along *ξ*_p_ carried out for at least 60 ns or 120 ns, respectively.

For PMFs along *ξ*_ch_, 24 umbrella positions were selected between 0.065 and 0.625 in steps of 0.08, and between 0.7 and 1.0 in steps of 0.02. For windows restrained to *ξ*_ch_ ≤ 0.7, an umbrella force constant of *k* = 3000 kJ mol^*−*1^ was used, and *k* = 5000 kJ mol^*−*1^ otherwise. For PMFs along *ξ*_p_, 63 umbrella positions were selected between 0.065 and 0.625 in steps of 0.08, between 0.68 and 1.10 in steps of 0.03, and between 1.12 and 6.97 in steps of 0.15. The following force constants were used: *k* = 3000 kJ mol^*−*1^ for *ξ*_p_ ≤ 0.75, *k* = 5000 kJ mol^*−*1^ for 0.75 ≤ *ξ*_p_ ≤ 1.1, and *k* = 400 kJ mol^*−*1^ otherwise. For systems containing no DMSO, each window was simulated for 100 ns, and the first 40 ns were omitted for equilibration. For systems containing DMSO, each window was simulated for 50 ns, and the first 10 ns were omitted for equilibration. During umbrella sampling, the pressure was controlled with the Parrinello-Rahman barostat (*τ* = 1 ps). All other parameters were chosen as described above. The PMFs were computed with the weighted histogram analysis method,^45^ as implemented in the gmx wham module of Gromacs.^46^ Statistical errors were estimated using 50 rounds of Bayesian bootstrapping of histograms.^46^ Here, each PMF computed from bootstrapped histograms was defined to zero at the PMF minimum, corresponding to the flat membrane, before computing the standard deviation between the bootstrapped PMFs. The thickness of the cylinder slices was set to *d_s_* = 0.1 nm. The degree to which a cylinder slice was considered as filled upon the addition of the first polar atom was taken as *ζ* = 0.75, as described previously.^32^ The number of slices *N_s_* was chosen such that the *ξ*_ch_ ≈ 0.2 referred to the flat membrane. This convention yields 26 and 32 slices for DMPC and POPC membranes, respectively.

To exclude that the integration time step or the algorithm used for temperature coupling influence the PMFs, we computed a PMF for pore nucleation along *ξ*_ch_ coordinate using two setups: (a) a 4 fs time step with a stochastic dynamics integration scheme^47^ and (b) a 5 fs time step and the temperature coupling indicated above. Because the PMFs were virtually identical, we used the computationally more efficient setup [option (b)].

## Results and Discussion

### Fixed versus mobile transmembrane cylinder

Before turning to the new reaction coordinate (RC), we investigated the influence of placing the transmembrane cylinder at a fixed lateral position, versus allowing the cylinder to follow the defect as the defect travels laterally in the membrane. As outlined in the Methods, the chain coordinate *ξ*_ch_ is defined by a transmembrane cylinder spanning the two polar head group regions and the hydrophobic membrane core. Using such a cylinder is critical because it avoids that two laterally displaced partial defects, one connected with the lower and one connected with the upper head groups, are misinterpreted as a continuous membrane-spanning defect. In other words, the cylinder enforces that the formed defect is localized in the membrane plane.

Previously, we found that it is critical to define the lateral position of the cylinder dynamically, allowing the cylinder to follow the defect. This protocol excludes that the system moves along the RC by shifting the defect laterally out of the cylinder, whereupon the nucleation barrier would be integrated out (Fig. 2, black curve). With a small simulation system of 128 lipids, simulations with such a mobile cylinder are numerically stable. However, upon simulating pore formation in large membrane patches, we found that using a mobile cylinder in umbrella windows with a nearly flat membrane may lead to integration problems, merely because water fluctuations at the glycerol region cause rapid lateral movements of the cylinder. To solve such numerical problems, we here fixed the cylinder in umbrella windows with reference position *ξ*_ch_ < 0.7, thereby avoiding integration problems, while leaving the cylinder mobile in windows with *ξ*_ch_ ≥ 0.7, thereby properly sampling the nucleation barrier. As shown in Fig. 2 (blue and magenta curves), fixing the cylinder at small *ξ*_ch_ leads to a slightly increased nucleation free energy by approx. 8 kJ/mol, rationalized by the entropic cost for forming the initial defect at a lateral position near the fixed cylinder center as compared to forming the defect at an arbitrary lateral position. However, the overall shape of the PMF remains similar. Hence, fixing the cylinder at small *ξ*_ch_ allowed numerically stable simulations of pore nucleation in both small and large membranes.

**Figure 2:**
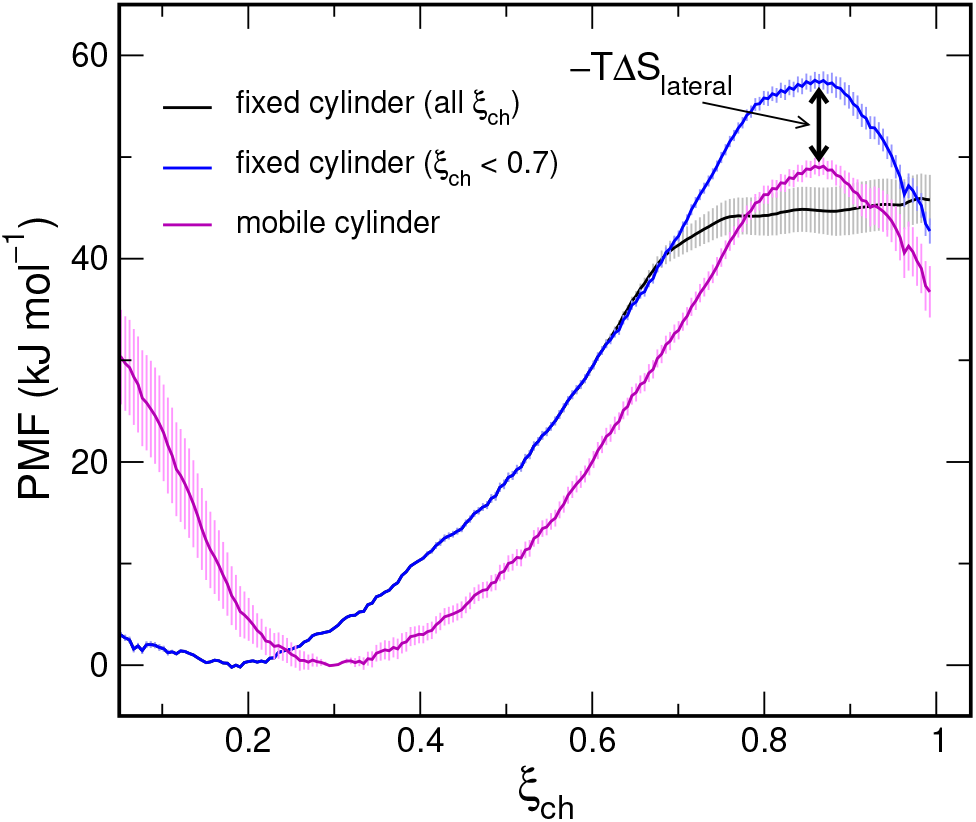
On the importance of using a mobile transmembrane cylinder: PMFs of pore nucleation over a membrane with 128 DMPC lipids, computed along the chain coordinate *ξ*_ch_. The PMF was computed with the transmembrane cylinder fixed in all umbrella windows, where the nucleation barrier was integrated out (black), with the cylinder fixed in none of the umbrella windows (magenta), or fixed only in windows whose umbrella center is <0.7 (blue). Shaded areas indicated standard errors computed by bootstrapping.

### A single PMF for nucleation and pore expansion

Figure 3 presents PMFs covering both pore nucleation and pore expansion across membranes of 512 DMPC or 512 POPC lipids, computed along the new RC *ξ*_p_. During umbrella sampling, we fixed the lateral position of the transmembrane cylinder in windows with a reference position of *ξ*_p_ < 0.7, and the cylinder was mobile in all other windows, equivalent to the setup used for Fig. 2 (blue curve). Typical simulation snapshots, taken from the final frames of various umbrella windows, reveal the progression from a flat membrane, via a partial defect with a locally thinned membrane due to indented head groups, a thin water needle, up to a fully formed large pore (Fig. 4A-H). The final snapshots of all umbrella windows are shown in Movie S1. In addition, to reveal the average shape of the defect at various *ξ*_p_ positions, Movie S2 presents the average densities of head groups, tails and water during all windows. Evidently, the PMFs reveal an approximately quadratic regime (*ξ*_p_ < 0.85), corresponding to the indentation of the head groups and penetration of a water needle into the membrane. Broad maxima in the PMFs around *ξ*_p_ ≈ 1 indicate the transition state of pore formation, followed by a shallow, broad minimum around *ξ*_p_ ≈ 2, indicating metastable (long-living) open pores. During further expansion of the pores, the PMFs reveal a linear regime at *ξ*_p_ > 4, which is compatible with a constant line tension of the pore rim.

**Figure 3:**
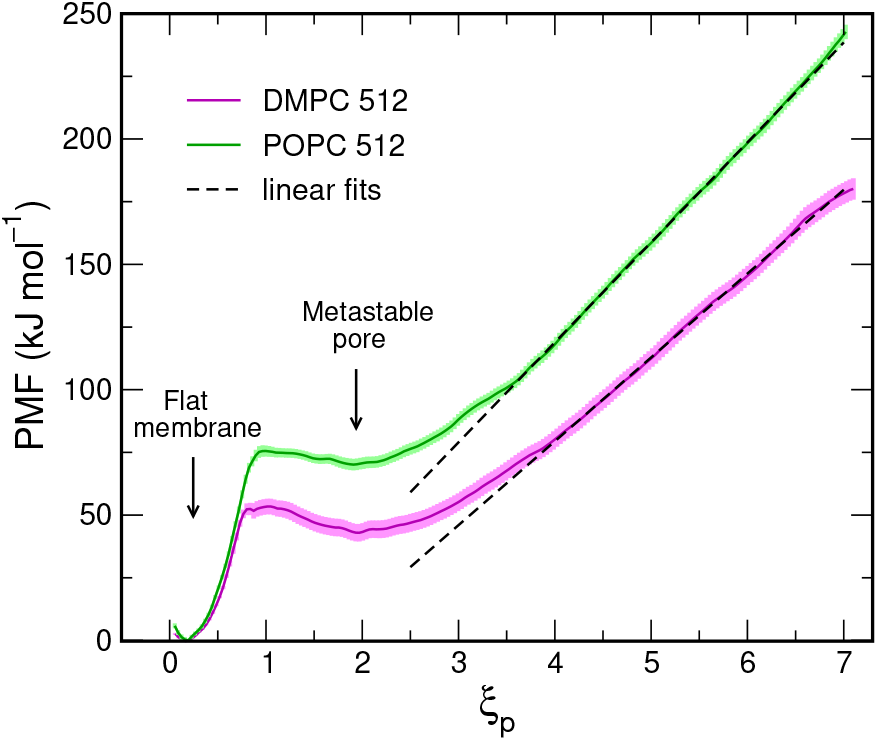
PMFs of pore nucleation and pore expansion in membranes of 512 DMPC lipids (magenta) or 512 POPC lipids, computed along the new reaction coordinate *ξ*_p_. The radii of the pores are given by *ξ*_p_*R*_0_, where *R*_0_ equals 0.405 nm and 0.380 for DMPC and POPC, respectively. Shaded areas indicate standard errors computed with bootstrapping.

**Figure 4:**
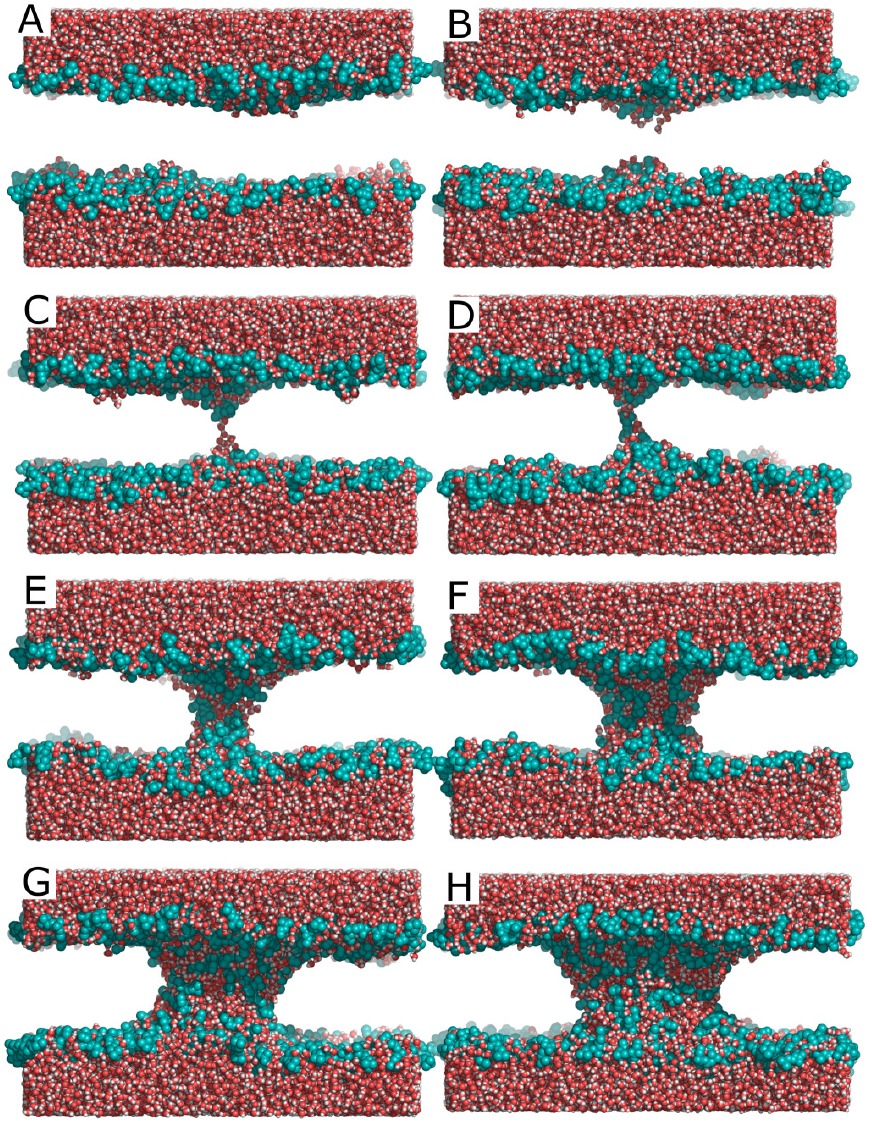
Typical snapshots during pore nucleation and expansion, taken from the final frames of umbrella sampling windows. The average *ξ*_p_ values of these windows were (from A–H) 0.09, 0.68, 0.84, 1.04, 2.16, 3.59, 5.08 and 6.56.

Comparing the PMFs for DMPC and POPC reveals a number of differences. First, the free energy of pore nucleation, given by the height of the barrier, is increased in POPC as compared to DMPC by 22 kJ/mol. This finding is compatible with previous studies that demonstrated that pore nucleation free energies correlate with membrane thickness. ^11,26^ Consequently, the free energy of the open pore is lower in DMPC than in POPC by 27 kJ/mol. Second, the depth of the shallow local minimum Δ*G*_cl_ at *ξ* ≈ 2 (relative to the barrier) is 10 kJ/mol for DMPC, but only 5 kJ/mol for POPC. Assuming that the rate of closing the pore is proportional to exp(−*β*Δ*G*_cl_), these values imply that the pore in DMPC has an ~8 times longer lifetime as compared to the pore in POPC. Structurally, this difference is rationalized by the larger spontaneous curvature of DMPC: with it larger headgroup-to-tail volume ratio, DMPC can better accommodate the large curvature at the pore rim. In contrast, the radius of the metastable pore is only marginally larger in DMPC as compared to POPC. The radius is given by *ξ*_p_*R*_0_ at the shallow PMF minimum, where the parameter *R*_0_ was 0.40 and 0.38 for DMPC and POPC, respectively (see Methods), implying radii of the metastable pores of 0.80 nm and 0.72 nm for DMPC and POPC, respectively.

Third, the slope of the PMF in the linear regime (*ξ*_p_ > 4) reveals the line tension of the pore edge. Classical nucleation theory models the free energy of the open pore by an unfavorable contribution from line tension of the pore edge plus a relief of surface tension between membrane and water (Eq. 1). The term owing to surface tension vanishes in our simulations because we simulated at constant number and not at constant chemical potential of lipids. Fitting a line to the PMFs in the interval *ξ*_p_ ∈ [4, 6.6] yields line tensions of (21.83 ± 0.04) pN and (27.69 ± 0.04) pN for DMPC and POPC, respectively. These lines tensions are slightly smaller than the values (23.3 ± 0.6) pN and (35.5 ± 0.05) pN for DMPC and POPC, respectively, reported by West *et al.*^48^ for pores in membranes modeled with the Berger lipids, as also used here. The slightly increased line tensions by West *et al.* might originate (i) from a different definition for the pore radius, or (ii) from simulating a smaller membrane patch of 256 lipids as compared to the patch of 512 lipids used here. In a previous study, we likewise observed that the line tension in a smaller DMPC system with 200 lipids steadily increased up to ~30 pN upon growing the pore.^11^ A detailed analysis of the effect of system size on the PMFs is presented below.

Taken together, with a single PMF along the new RC, we obtain the full free energy landscape of pore nucleation and pore expansion. The PMF displays the free energy cost for nucleating the pore, the metastability, size, and free energy of the open pore, as well as the line tension of the pore edge.

### Switching the reaction coordinate from nucleation to expansion

To illustrate how the RC *ξ*_p_ accounts for the switch from pore nucleation to pore expansion, Fig. 5 presents the progression of the most relevant parameters during nucleation and expansions. The values in Fig. 5 report the respective mean and standard deviation taken from umbrella windows of the system with 512 POPC lipids. Evidently, during pore nucleation (*ξ*_p_ ≲ 0.9), the new RC *ξ*_p_ (red symbols) is simply given by the chain coordinate *ξ*_ch_ (black symbols) because the switch function *H_ϵ_* (green symbols) still takes the value of zero (see also Eq. 4). As the chain coordinate approaches the switch value 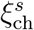 set to 0.925 is this simulation, the smoothed Heaviside step function 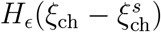 switches from zero to unity and, hence, the new RC *ξ*_p_ starts to deviate from the chain coordinate *ξ*_ch_ (see Eq. 4, and Fig. 5, black/red/green symbols). Henceforth, *ξ*_ch_ saturates at unity (indicating a continuous water defect), whereas *ξ*_p_ continues to increase as (*R* − *R*_0_)*/R*_0_ (orange symbols).

**Figure 5:**
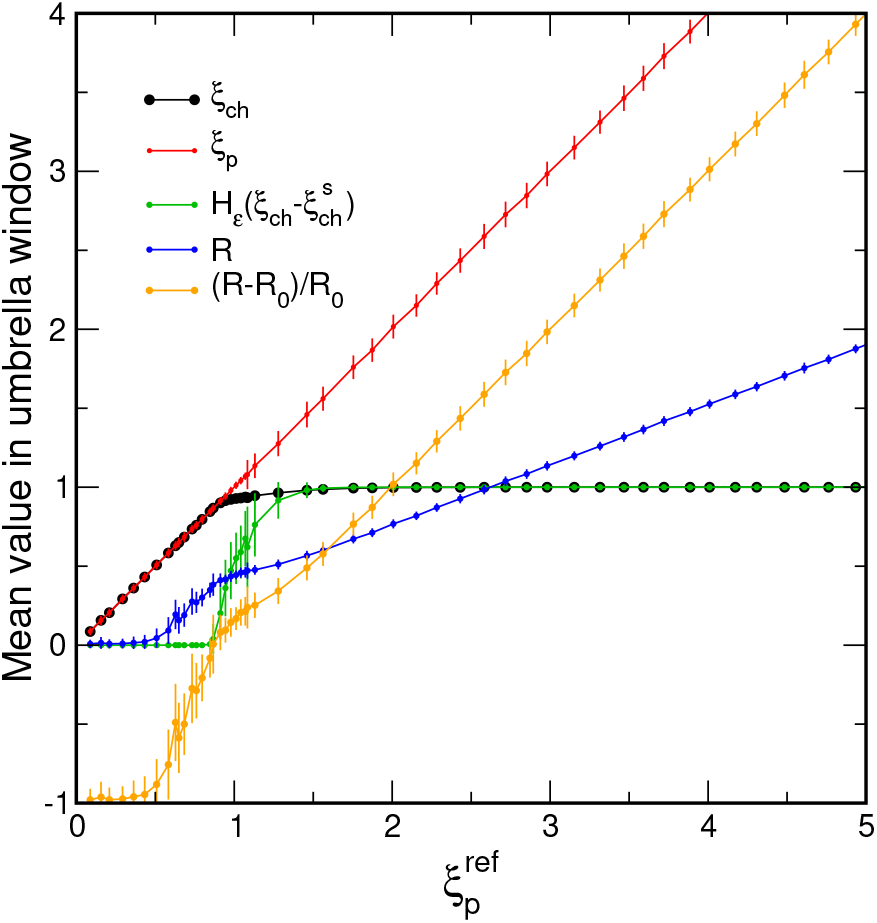
Mean values of *ξ*_p_, the chain coordinate *ξ*_ch_, the switch function *H_ϵ_*, pore radius *R*, and (*R* − *R*_0_)*/R*_0_, taken from umbrella sampling windows with reference position 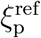 of the system with 512 POPC lipids. Vertical bars indicate the standard deviation within the windows.

In addition, Fig. 5 demonstrates the relation between the pore radius *R* and *ξ*_p_. During pore nucleation (0 < *ξ*_p_ ≲ 0.9), *R* increases up to the unit radius *R*_0_, corresponding to the radius of a continuous thin polar defect, where *R*_0_ = 0.38 in this simulation (blue symbols). Henceforth, during pore growth, *R* increases linearly as *ξ*_p_*R*_0_. Overall, Fig. 5 illustrates how a restraint along *ξ*_ch_ is automatically turned into a restraint along *R*, thereby driving pore nucleation and expansion by restraining only a single degree of freedom.

### The simulation box size influences the size and the free energy of the open pore

Previous simulations investigated pore formation in membrane of various size, spanning systems between 64 and 512 lipids.^17,20,24–26,28,31,32,49–52^ However, we previously found that the system size may strongly influence the free energy landscape of pore formation. In small membrane systems, the open pore may be destabilized because there is no flat membrane between the pore and its periodic image.^31^

To probe the influence of system size on the free energy landscape of pore formation, we computed the PMF along the newly proposed RC using systems of 128, 200, or 512 DMPC lipids. As shown in Fig. 6, the free energy of pore nucleation, as indicated by the local maximum at *ξ*_p_ ≈ 0.9, is not influenced by system size. In contrast, in the regime of expansion (*ξ*_p_ > 1.5), major effects of system size are apparent. As expected, pores are destabilized in smaller systems due to interactions of the pore with its periodic image. Specifically, with 128 DMPC lipids, the free energy of the metastable pore is increased as compared to larger membranes, suggesting a reduced lifetime of the pore (Fig. 6, black). In addition, the local minimum of the PMF is shifted to smaller *ξ*_p_, demonstrating that the metastable pore with 128 DMPC lipids has a reduced radius.

**Figure 6:**
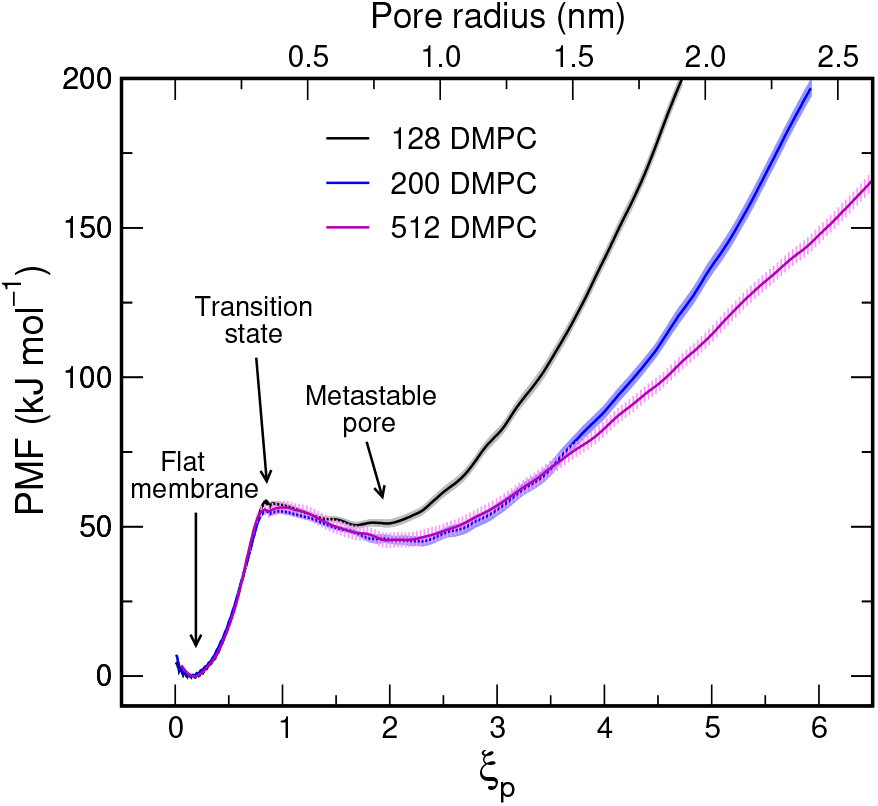
PMFs of pore nucleation and expansion in DMPC membranes with 128 (black), 200 (blue), or 512 DMPC lipids (magenta), shown as a function of *ξ*_p_ (lower abscissa). The upper abscissa shows the pore radius, computed as *ξ*_p_*R*_0_ where *R*_0_ = 0.405 nm.

The PMFs for 200 or 512 DMPC lipids, in contrast, agree around the minimum of the metastable pore, suggesting that a 200 DMPC system is sufficient for capturing the energetics and radius of the metastable state (Fig. 6, blue and magenta). However, upon further expansion of the pore (*ξ*_p_ > 3.5), the PMF for 512 lipids reveals a constant slope indicating a constant line tension (see above), whereas the slope of the PMF for 200 lipids continuously increases. This behavior suggest that the 200 lipid system cannot be used to compute the line tension of an expanding pore because the system passes directly from (i) the regime with reduced (or even zero) line tension around the metastable state to (ii) a regime with spuriously increased line tension due to periodic boundary artifacts.

### Case study: Effect of DMSO on the PMF of pore nucleation and expansion

Dimethyl sulfoxide (DMSO) is a widely used organic solvent, whose effect on lipid membranes has been studied in great detail by experiments and simulation. DMSO influences the structure, hydration properties, phase behavior, and permeability of membranes.^53–61^ MD simulations showed that DMSO preferentially binds at the interface between the polar and apolar regions of the membrane, where it may act as a spacer, thereby rendering the membrane thinner and floppier.^62^ At high DMSO concentrations, the formation of water defects has been observed in unbiased simulation studies, rationalizing the increased permeability of membranes for electrolytes in presence of DMSO.^62–65^ However, the effects of the DMSO on the free energies of pore formation and expansion are not well understood. Here, we revisit DMSO to illustrate how the proposed RC may be used to rationalize the effects of DMSO on membrane stability and pore formation in energetic terms.

Figure 7 presents PMFs of pore nucleation and expansion in a membrane with 512 POPC membrane plus increasing amount DMSO in POPC:DMSO ratios of 2:1, 1:1, 1:1.5, and 1:2. Evidently, the addition of DMSO has multiple, drastic effects on the PMF. (i) The pore nucleation free energy decreases by up to 20 kJ/mol, indicating that the rate of pore opening is increased by a factor of ~3000; (ii) The free energy of the open, metastable pore decreases by up to 40 kJ/mol, demonstrating a greatly increased probability for a open pore. (iii) As a consequence of points (i) and (ii), the free energy barrier that must be overcome to close the pore increases with DMSO, implying greatly decreased rate of pore closing (or increased pore lifetime) by a factor of ~3000. Hence, the pore becomes more metastable. (iv) The local PMF minimum corresponding to the open, metastable pore is shifted to larger *ξ*_p_, implying larger pore radii. However, because the minima are shallow, the pores likely adopt a wide distribution of pore radii. (v) The slope of the PMFs at large *ξ*_p_ decreases, reflecting a decreased line tension of the pore rim (Fig. 7, dashed lines). The line tensions obtained from the slope of the PMFs are summarized in Table 1. The larger radii of the metastable pores at increased DMSO content (point iv) are likely a consequence of the smaller line tension. In addition, the reduced line tension affects the shape of large open pores. Whereas pores adopt approximately circular shapes at low DMSO content (Fig. 8A–C), as favored by a high line tension of the rim, pores at high DMSO content adopt increasingly irregular shapes (Fig. 8D–E). This trend was confirmed by visual inspection of many more MD frames. (vi) Finally, the minima at *ξ*_p_ ≈ 0.3 corresponding to the flat, unperturbed membrane are shifted to slightly larger *ξ*_p_. This shift reflects that, in the flat membrane, more slices of the transmembrane cylinder (Fig. 1A) are filled by polar atoms; hence, the flat membrane is thinner in presence of DMSO. ^62^

**Figure 7:**
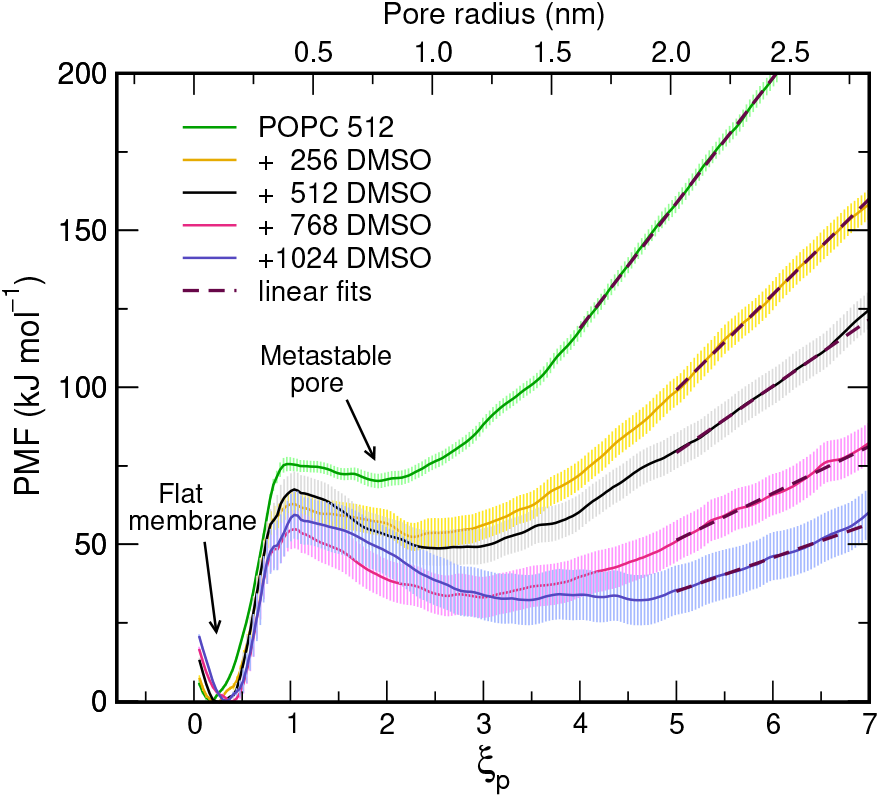
PMFs of pore nucleation and expansion in membranes of 512 POPC lipids plus an increasing number of DMSO molecules (see legend). The upper abscissa shows the pore radius, computed as *ξ*_p_*R*_0_ where *R*_0_ = 0.405 nm.

**Figure 8:**
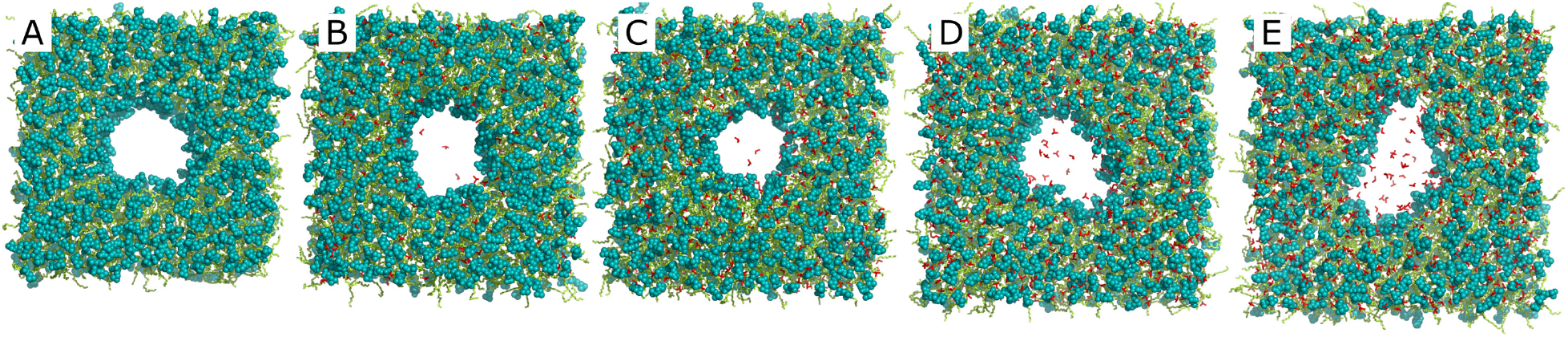
Simulation snapshots of membranes of 512 POPC lipids with a large pore, containing (A) no DMSO or an increasing content of (B–E) 256, 512, 768, 1024 DMSO molecules, respectively. Snapshots were taken from the final frames of the umbrella sampling window restrained to *ξ*_p_ 0.7. Headgroups are shown as cyan spheres, tails as lime sticks, and DMSO as red sticks. Water is not shown for clarity.

**Table 1:** Line tension of the rim of large pores over a POPC membrane in absence or presence of DMSO, obtained from the slope of the PMFs at large pore radii (Fig. 7, dashed lines).

| Composition | Line tension (pN) |
| --- | --- |
| 512 POPC | 27.7 |
| +256 DMSO | 12.7 |
| +512 DMSO | 8.9 |
| +768 DMSO | 6.2 |
| +1024 DMSO | 4.5 |

Taken together, DMSO shows a multitude of effects on the free energy landscape of pore formation, affecting the rates of pore nucleation and closure, the probability and size of the pore size, as well as the free energy cost for further expansion. The PMFs along *ξ*_p_ reflect all these effects, hence providing comprehensive understanding of the influence of DMSO on the energetics of membrane stability and pore formation.

## Discussion

We have presented a new reaction coordinate (RC) *ξ*_p_ for pore formation by combining two established RCs, namely (i) the “chain coordinate” *ξ*_ch_ to follow pore nucleation, and (ii) the pore radius *R* to follow pore expansion. This was achieved by introducing a continuous switch from one coordinate to the other. Because pulling the system along the new RC *ξ*_p_ allows for reversible pathways from the flat, unperturbed membrane up to a large pore with a diameter of several nanometers, the RC allows one to compute the free energy of the pore (at any stage of pore formation) relative to the flat, unperturbed membrane.

The capability of new RC was illustrated by computing the PMFs of pore formation with different lipids and with different system sizes. For DMPC membranes, a system of 128 lipids was sufficient for obtaining the correct barrier for pore nucleation, 200 lipid were required to obtain the correct free energy and size of a metastable pore, but 512 lipids were required for reliably obtaining the line tension of the pore rim. Hence, the appropriate system size needed to avoid periodic boundary artifacts depends on the quantity under study. In addition, we showed that DMSO has drastic effects on pore formation, affecting the free energy barrier of pore nucleation, the free energy and metastability of the open pore, and the line tension as given by the slope of the PMF at large pore radii.

Pore nucleation and pore expansion occur along different directions in conformation space. Therefore, it may appear surprising that restraining only a single degree of freedom, as given by *ξ*_p_, is sufficient for steering the system along such different directions. However, the success of the RC used here critically depends on the metastability of the pore. After pore nucleation, and as the radius of the pore starts to increase, the metastability ensures that the transmembrane defect remains intact, that the chain coordinate *ξ*_ch_(**r**) remains close to unity and, consequently, that also the switch function 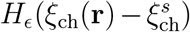 remains close to unity. Therefore, while restraining the open pore along *ξ*_p_, the switch function *H_ϵ_* does not fall back to small values, and we obtain a stable restraint along the radius *R*. As such, the design of the new RC makes use of the specific free energy landscape of the pore formation. We anticipate that PMF calculations with highly unstable pores alternative strategies might be advantageous. For instance, expanding the pore with an repulsive cylindrical potential acting on the lipid tail atoms might be less prone to undesired effects as compared the attractive potential acting on polar atoms used here.

The molecular mechanism of pore formation observed here is largely compatible with previous studies, suggesting that pore formation by pulling along *ξ*_p_ does not perturb the membrane more strongly than needed for pore formation. In other words, this suggests that is the simulations follow approximately the minimum free energy pathway. As visualized in Movies S1 and S2 and by Fig. 4, pore formation starts by a local thinning of the membrane, that is by an elastic indentation of the headgroup/tail interface. ^14^ Next, a thin water needle penetrates the hydrophobic core.^26^ The water needle widens by penetration of additional water and a substantial number of lipid headgroups. Interestingly, the density analysis in Movie S2 reveals that small pores with a radius of 4–7Å contain more headgroup than water density. This observation is compatible with an important role of the headgroups for crossing the transition state of pore formation, as suggested by a recent study.^14^ Larger pores with a radius of more than 12 Å reveal a nearly ideal toroidal shape, in which the head groups shield the membrane-spanning water defect from the hydrophobic lipid tails. However, the density of the headgroups is lower in the toroidal pore as compared to the flat membrane, reflecting a sub-optimal lipid packing at the pore rim (Movie S2).

The RC proposed here will be useful for a range of applications. It will be interesting to quantify the influence of antimicrobial or cell-penetrating peptides on pore formation, as required for a rational design of such membrane-active peptides. ^66^ PMF calculations with the new RC could reveal the free energy landscape of electroporation. ^20^ The RC may be further applied to quantify the stability of liposome-based drug carries, and to model the controlled drug release from such carriers.^67^ Routine calculations of the pore’s line tension may be useful for further refinement of lipid force fields. Indeed, agreement of lipid membrane simulations with experimental line tension data would be important for simulations that involve topological transitions of membranes, during which membranes adopt non-planar geometries. In the context of membrane fusion, PMFs could rationalize the role of the lipid composition and of transmembrane helices during fusion pore opening and expansion.^68^ In addition, we anticipate that the RC may be used to study stalk formation and stalk widening, during which the hydrophobic membrane cores carry out similar topological transition as the water phase during pore formation and expansion.^3,69^

## Appendix

### Gradients of *ξ_p_(r)*

Restraining the simulation system along *ξ*_p_(**r**), as carried out during umbrella sampling simulations, requires the calculation of the gradient ∇_*i*_*ξ*_p_ of the RC with respect to the Cartesian coordinates of atom *i*. From Eq. 4, we have

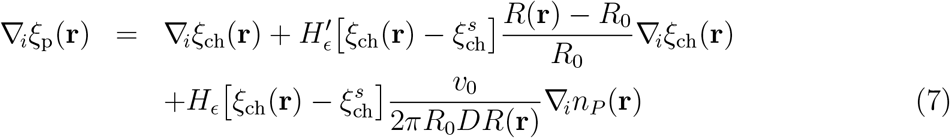

Here, 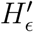 is the derivative of the smoothed Heaviside step function. The gradient of the chain coordinate ∇_*i*_*ξ*_ch_ was computed as described previously.^32^ The last term of Eq. 7 was set to zero if *R*(**r**) = 0. The gradient of the number of polar atoms is

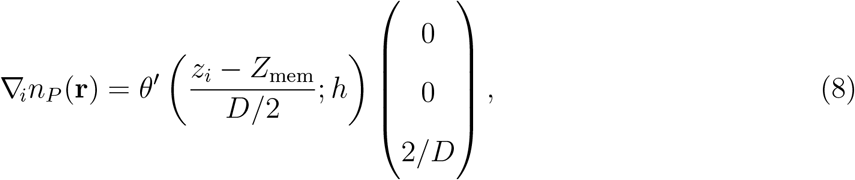

where *θ*′ is the derivative of *θ*.

### Modification of the RC and simulation setup for highly floppy membranes

By default, the radius of the pore was defined based on the number of polar atoms within a horizontal slab at the membrane center (see Methods and Fig. 1B, orange shading). For large, floppy membranes, such as membranes with 512 lipids in presence of DMSO, this definition leads to problems because polar atoms may enter the slab not only by expanding the pore but also by membrane undulations. In such cases, we replaced the horizontal slab with a cylinder. The axis of this cylinder coincided with the axis of the membrane-spanning cylinder used for defining the chain coordinate *ξ*_ch_. The radius was chosen 0.8 nm larger than the current approximate radius of the pore given by 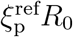, where 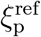 is the reference position of the umbrella potential along *ξ*_p_. This definition reduced undesired effects from membrane undulations, while at the same time minimized undesired effects of the cylinder on shape fluctuations of the pore.

For the floppiest membranes (512 POPC plus 768/1024 DMSO), an additional measure was needed to avoid problems from heavy membrane undulations. Here, we used flat-bottomed position restraints on *z*-coordinates all lipid tails atoms, *V*_fb_(*z_i_*) = *k/*2(|*z_i_* − *Z*_mem_| − *z*_fb_)^2^*H*[|*z_i_* − *Z*_mem_| − *z*_fb_]. Here *z_i_* is the *z*-coordinate (membrane normal) of tail atom *i*, *Z*_mem_ the center of the membrane, *z*_fb_ = 2.2 nm the half-width of the flat region, *k* = 100 kJ/mol/nm^2^ the force constant, and *H* is the Heaviside step function. This potential allowed natural local fluctuations of head groups but excluded large-scale membrane undulations, thereby rendering the simulations more stable.

## Acknowledgement

I thank Greg Bubnis for stimulating discussions and for critically reading the manuscript. This study was supported by the Deutsche Forschungsgemeinschaft via grant SFB 1027/B7.

## Supporting Information Available

The following files are available free of charge.

- Movie S1: 62 simulation snapshots of a membrane of 512 POPC lipids during pore formation, taken from the final MD frames of umbrella sampling windows.
- Movie S2: density of head groups, water, and lipid tails, averaged over each of 62 umbrella windows of a membrane of 512 POPC lipids. The average *ξ*_p_ and the radius *R* of pore are shown in the top right corner. The density is plotted in color code as function of the lateral distance *r* and vertical distance *z* from the pore center.

## Graphical TOC Entry

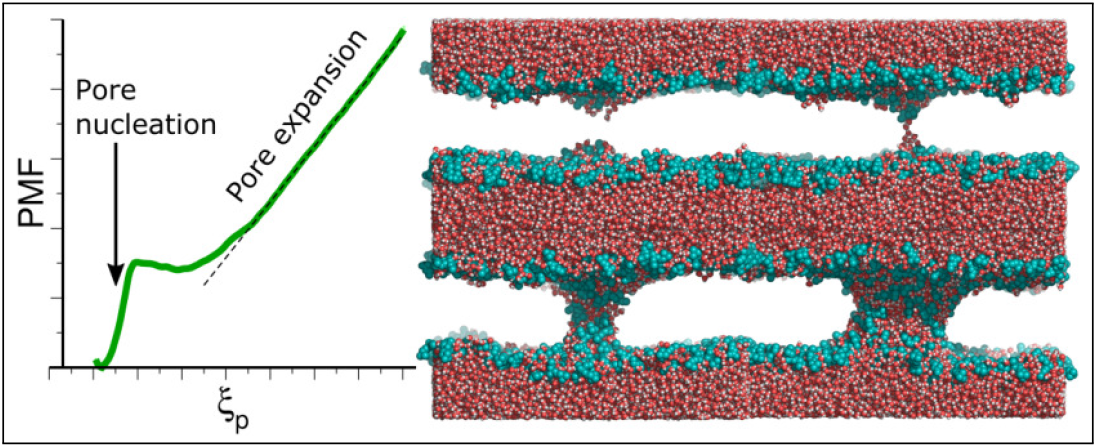

## References

(1) Brogden, K. A. Antimicrobial peptides: pore formers or metabolic inhibitors in bacteria? Nat Rev Microbiol 2005, 3, 238–250.

(2) Peraro, M. D.; van der Goot, F. G. Pore-forming toxins: ancient, but never really out of fashion. Nat Rev Microbiol 2016, 14, 77–92.

(3) Jahn, R.; Lang, T.; Südhof, T. C. Membrane fusion. Cell 2003, 112, 519–533.

(4) Risselada, H. J.; Grubmüller, H. How SNARE molecules mediate membrane fusion: recent insights from molecular simulations. Curr. Opin. Struct. Biol. 2012, 22, 187–196.

(5) Majd, S.; Yusko, E. C.; Billeh, Y. N.; Macrae, M. X.; Yang, J.; Mayer, M. Applications of biological pores in nanomedicine, sensing, and nanoelectronics. Current Opinion in Biotechnology 2010, 21, 439–476.

(6) Abidor, I.; Arakelyan, V.; Chernomordik, L.; Chizmadzhev, Y. A.; Pastushenko, V.; Tarasevich, M. Electric breakdown of bilayer lipid membranes I. The main experimental facts and their qualitative discussion. Bioelectrochem. Bioenerg. 1979, 6, 37–51.

(7) Kotnik, T.; Frey, W.; Sack, M.; Haberl Meglič, S.; Peterka, M.; Miklavčič, D. Electroporation-based applications in biotechnology. Trends in Biotechnology 2015, 33, 480–488.

(8) Zhelev, D. V.; Needham, D. Tension-stabilized pores in giant vesicles: determination of pore size and pore line tension. Biochimica et Biophysica Acta (BBA) - Biomembranes 1993, 1147, 89–104.

(9) Levin, Y.; Idiart, M. A. Pore dynamics of osmotically stressed vesicles. Physica A: Statistical Mechanics and its Applications 2004, 331, 571–578.

(10) Chabanon, M.; Ho, J. C.; Liedberg, B.; Parikh, A. N.; Rangamani, P. Pulsatile Lipid Vesicles under Osmotic Stress. Biophys. J. 2017, 112, 1682–1691.

(11) Ting, C. L.; Awasthi, N.; Müller, M.; Hub, J. S. Metastable Prepores in Tension-Free Lipid Bilayers. Phys. Rev. Lett. 2018, 120.

(12) Bennett, W. D.; Tieleman, D. P. The Importance of Membrane Defects Lessons from Simulations. Acc. Chem. Res. 2014, 47, 2244–2251.

(13) Grafmüller, A.; Knecht, V. The free energy of nanopores in tense membranes. Phys. Chem. Chem. Phys. 2014, 16, 11270.

(14) Bubnis, G.; Grubmüller, H. Sequential water and headgroup merger: Membrane poration paths and energetics from MD simulations. BioRxiv 2020, doi: 10.1101/2020.06.15.152215.

(15) Taupin, C.; Dvolaitzky, M.; Sauterey, C. Osmotic pressure-induced pores in phospholipid vesicles. Biochemistry 1975, 14, 4771–4775.

(16) Litster, J. Stability of lipid bilayers and red blood cell membranes. Physics Letters A 1975, 53, 193–194.

(17) Tieleman, D. P.; Leontiadou, H.; Mark, A. E.; Marrink, S.-J. Simulation of pore formation in lipid bilayers by mechanical stress and electric fields. J. Am. Chem. Soc. 2003, 125, 6382–6383.

(18) Leontiadou, H.; Mark, A. E.; Marrink, S. J. Molecular dynamics simulations of hydrophilic pores in lipid bilayers. Biophys. J. 2004, 86, 2156–2164.

(19) Tarek, M. Membrane electroporation: a molecular dynamics simulation. Biophys. J. 2005, 88, 4045–4053.

(20) Böckmann, R. A.; De Groot, B. L.; Kakorin, S.; Neumann, E.; Grubmüller, H. Kinetics, statistics, and energetics of lipid membrane electroporation studied by molecular dynamics simulations. Biophys. J. 2008, 95, 1837–1850.

(21) Piggot, T. J.; Holdbrook, D. A.; Khalid, S. Electroporation of the E. coli and S. aureus membranes: molecular dynamics simulations of complex bacterial membranes. J. Phys. Chem. B 2011, 115, 13381–13388.

(22) Wang, Z.-J.; Frenkel, D. Pore nucleation in mechanically stretched bilayer membranes. J. Chem. Phys. 2005, 123, 154701.

(23) Dixit, M.; Lazaridis, T. Free energy of hydrophilic and hydrophobic pores in lipid bilayers by free energy perturbation of a restraint. J. Chem. Phys. 2020, 153, 054101.

(24) Wohlert, J.; Den Otter, W.; Edholm, O.; Briels, W. Free energy of a trans-membrane pore calculated from atomistic molecular dynamics simulations. J. Chem. Phys. 2006, 124, 154905.

(25) Tieleman, D. P.; Marrink, S.-J. Lipids out of equilibrium: Energetics of desorption and pore mediated flip-flop. J. Am. Chem. Soc. 2006, 128, 12462–12467.

(26) Bennett, W. D.; Sapay, N.; Tieleman, D. P. Atomistic simulations of pore formation and closure in lipid bilayers. Biophys. J. 2014, 106, 210–219.

(27) Sun, D.; Forsman, J.; Woodward, C. E. Atomistic Molecular Simulations Suggest a Kinetic Model for Membrane Translocation by Arginine-Rich Peptides. J. Phys. Chem. B 2015, 119, 14413–14420.

(28) Mirjalili, V.; Feig, M. Density-Biased Sampling: A Robust Computational Method for Studying Pore Formation in Membranes. J. Chem. Theory Comput. 2014, 11, 343–350.

(29) Akimov, S. A.; Volynsky, P. E.; Galimzyanov, T. R.; Kuzmin, P. I.; Pavlov, K. V.; Batishchev, O. V. Pore formation in lipid membrane II: Energy landscape under external stress. Sci. Rep. 2017, 7, 12509.

(30) Akimov, S. A.; Volynsky, P. E.; Galimzyanov, T. R.; Kuzmin, P. I.; Pavlov, K. V.; Batishchev, O. V. Pore formation in lipid membrane I: Continuous reversible trajectory from intact bilayer through hydrophobic defect to transversal pore. Sci. Rep. 2017, 7.

(31) Awasthi, N.; Hub, J. S. Simulations of pore formation in lipid membranes: reaction coordinates, convergence, hysteresis, and finite-size effects. J. Chem. Theory Comput. 2016, 12, 3261–3269.

(32) Hub, J. S.; Awasthi, N. Probing a continuous polar defect: A reaction coordinate for pore formation in lipid membranes. J. Chem. Theory Comput. 2017, 13, 2352–2366.

(33) Abraham, M. J.; Murtola, T.; Schulz, R.; Páll, S.; Smith, J. C.; Hess, B.; Lindahl, E. GROMACS: High performance molecular simulations through multi-level parallelism from laptops to supercomputers. SoftwareX 2015, 1, 19–25.

(34) Knight, C. J.; Hub, J. S. MemGen: A general web server for the setup of lipid membrane simulation systems. Bioinformatics 2015, 31, 2897–2899.

(35) Berger, O.; Edholm, O.; Jähnig, F. Molecular dynamics simulations of a fluid bilayer of dipalmitoylphosphatidylcholine at full hydration, constant pressure, and constant temperature. Biophys. J. 1997, 72, 2002–2013.

(36) Berendsen, H. J. C.; Postma, J. P. M.; Gunsteren, W. F. v.; Hermans, J. In Intermolecular Forces; Pullman, B., Ed.; D. Reidel Publishing Company: Dordrecht, 1981; pp 331–342.

(37) Geerke, D. P.; Oostenbrink, C.; van der Vegt, N. F. A.; van Gunsteren, W. F. An Effective Force Field for Molecular Dynamics Simulations of Dimethyl Sulfoxide and Dimethyl Sulfoxide-Water Mixtures. J. Phys. Chem. B 2004, 108, 1436–1445.

(38) Miyamoto, S.; Kollman, P. A. SETTLE: An Analytical Version of the SHAKE and RATTLE Algorithms for Rigid Water Models. J. Comp. Chem. 1992, 13, 952–962.

(39) Hess, B. P-LINCS: A Parallel Linear Constraint Solver for Molecular Simulation. J. Chem. Theory Comput. 2008, 4, 116–122.

(40) Bussi, G.; Donadio, D.; Parrinello, M. Canonical sampling through velocity rescaling. J. Chem. Phys. 2007, 126, 014101.

(41) Berendsen, H. J. C.; Postma, J. P. M.; DiNola, A.; Haak, J. R. Molecular dynamics with coupling to an external bath. J. Chem. Phys. 1984, 81, 3684–3690.

(42) Darden, T.; York, D.; Pedersen, L. Particle mesh Ewald: an N$\cdot$log(N) method for Ewald sums in large systems. J. Chem. Phys. 1993, 98, 10089–10092.

(43) Essmann, U.; Perera, L.; Berkowitz, M. L.; Darden, T.; Lee, H.; Pedersen, L. G. A smooth particle mesh ewald potential. J. Chem. Phys. 1995, 103, 8577–8592.

(44) Torrie, G. M.; Valleau, J. P. Monte Carlo free energy estimates using non-Boltzmann sampling: Application to the sub-critical Lennard-Jones fluid. Chem. Phys. Lett. 1974, 28, 578–581.

(45) Kumar, S.; Bouzida, D.; Swendsen, R. H.; Kollman, P. A.; Rosenberg, J. M. The weighted histogram analysis method for free-energy calculations on biomolecules. I. The method. J. Comput. Chem. 1992, 13, 1011–1021.

(46) Hub, J. S.; Groot, B. L. d.; Spoel, D. v. d. g_wham–A Free Weighted Histogram Analysis Implementation Including Robust Error and Autocorrelation Estimates. J. Chem. Theory Comput. 2010, 6, 3713–3720.

(47) Gunsteren, W. F. v.; Berendsen, H. J. C. A Leap-Frog Algorithm for Stochastic Dynamics. Mol. Sim. 1988, 1, 173–185.

(48) West, A.; Ma, K.; Chung, J. L.; Kindt, J. T. Simulation Studies of Structure and Edge Tension of Lipid Bilayer Edges: Effects of Tail Structure and Force-Field. J. Phys. Chem. A 2013, 117, 7114–7123.

(49) Bennett, W. D.; Tieleman, D. P. Water defect and pore formation in atomistic and coarse-grained lipid membranes: pushing the limits of coarse graining. J. Chem. Theory Comput. 2011, 7, 2981–2988.

(50) Huang, K.; García, A. E. Effects of truncating van der Waals interactions in lipid bilayer simulations. J. Chem. Phys. 2014, 141, 105101.

(51) Hu, Y.; Sinha, S. K.; Patel, S. Investigating hydrophilic pores in model lipid bilayers using molecular simulations: correlating bilayer properties with pore-formation thermodynamics. Langmuir 2015, 31, 6615–6631.

(52) Awasthi, N.; Kopec, W.; Wilkosz, N.; Jamróz, D.; Hub, J. S.; Zatorska, M.; Petka, R.; Nowakowska, M.; Kepczynski, M. Molecular Mechanism of Polycation-Induced Pore Formation in Biomembranes. ACS Biomater. Sci. Eng. 2019, 5, 780–794.

(53) Bunch, W. H. The effect of DMSO on the permeation of non-electrolytes through the barnacle cell membrane. J. Cell. Physiol. 1968, 72, 49–54.

(54) Gordeliy, V.; Kiselev, M.; Lesieur, P.; Pole, A.; Teixeira, J. Lipid Membrane Structure and Interactions in Dimethyl Sulfoxide/Water Mixtures. Biophys. J. 1998, 75, 2343–2351.

(55) Yu, Z.-W.; Quinn, P. J. The modulation of membrane structure and stability by dimethyl sulphoxide (Review). Mol. Membr. Biol. 1998, 15, 59–68.

(56) Kiselev, M.; Lesieur, P.; Kisselev, A.; Grabielle-Madelmond, C.; Ollivon, M. DMSO-induced dehydration of DPPC membranes studied by X-ray diffraction, small-angle neutron scattering, and calorimetry. J. Alloys Compd. 1999, 286, 195–202.

(57) Yamashita, Y.; Kinoshita, K.; Yamazaki, M. Low concentration of DMSO stabilizes the bilayer gel phase rather than the interdigitated gel phase in dihexadecylphosphatidylcholine membrane. Biochimica et Biophysica Acta (BBA) - Biomembranes 2000, 1467, 395–405.

(58) Smondyrev, A. M.; Berkowitz, M. L. Molecular Dynamics Simulation of DPPC Bilayer in DMSO. Biophys. J. 1999, 76, 2472–2478.

(59) Sum, A. K.; de Pablo, J. J. Molecular Simulation Study on the Influence of Dimethyl-sulfoxide on the Structure of Phospholipid Bilayers. Biophys. J. 2003, 85, 3636–3645.

(60) Notman, R.; den Otter, W. K.; Noro, M. G.; Briels, W.; Anwar, J. The Permeability Enhancing Mechanism of DMSO in Ceramide Bilayers Simulated by Molecular Dynamics. Biophys. J. 2007, 93, 2056–2068.

(61) Lee, Y.; Pincus, P. A.; Hyeon, C. Effects of Dimethyl Sulfoxide on Surface Water near Phospholipid Bilayers. Biophys. J. 2016, 111, 2481–2491.

(62) Notman, R.; Noro, M.; O’Malley, B.; Anwar, J. Molecular Basis for Dimethylsulfoxide (DMSO) Action on Lipid Membranes. J. Am. Chem. Soc. 2006, 128, 13982–13983.

(63) Gurtovenko, A. A.; Anwar, J. Modulating the Structure and Properties of Cell Membranes: The Molecular Mechanism of Action of Dimethyl Sulfoxide. J. Phys. Chem. B 2007, 111, 10453–10460.

(64) Hughes, Z. E.; Mark, A. E.; Mancera, R. L. Molecular Dynamics Simulations of the Interactions of DMSO with DPPC and DOPC Phospholipid Membranes. J. Phys. Chem. B 2012, 116, 11911–11923.

(65) de Ménorval, M.-A.; Mir, L. M.; Fernández, M. L.; Reigada, R. Effects of Dimethyl Sulfoxide in Cholesterol-Containing Lipid Membranes: A Comparative Study of Experiments In Silico and with Cells. PLoS ONE 2012, 7, e41733.

(66) Fjell, C. D.; Hiss, J. A.; Hancock, R. E.; Schneider, G. Designing antimicrobial peptides: form follows function. Nat. Rev. Drug Discovery 2012, 11, 37–51.

(67) Torchilin, V. P. Recent advances with liposomes as pharmaceutical carriers. Nat Rev Drug Discov 2005, 4, 145–160.

(68) Dhara, M.; Mantero Martinez, M.; Makke, M.; Schwarz, Y.; Mohrmann, R.; Bruns, D. Synergistic actions of v-SNARE transmembrane domains and membrane-curvature modifying lipids in neurotransmitter release. eLife 2020, 9, e55152.

(69) Smirnova, Y. G.; Risselada, H. J.; Müller, M. Thermodynamically reversible paths of the first fusion intermediate reveal an important role for membrane anchors of fusion proteins. Proc Natl Acad Sci USA 2019, 116, 2571–2576.

